# Regulation of osteoblast autophagy based on PI3K/AKT/mTOR signaling pathway study on the effect of β-ecdysterone on fracture healing

**DOI:** 10.1101/2021.05.28.446186

**Authors:** Yanghua Tang, Yafeng Mo, Dawei Xin, Zhenfei Xiong, Linru Zeng, Gan Luo

## Abstract

**Purpose:** To investigate the effects of β-ecdysterone on fracture healing and the underlying mechanism.

**Methods:** MTT assay was used to detect the cell viability and alkaline phosphatase (ALP) activity was measured using a commercial kit. AO/PI and flow cytometry assays were used to determine the state of apoptosis of osteoblasts. The expression level of RunX2, ATG7 and LC3 was evaluated by qRT-PCR and Western blot assays. X-ray and HE staining were conducted on the fractured femur to evaluate the pathological state. Immunohistochemical assay was used to detect the expression level of Beclin-1 and immunofluorescence assay was used to measure the expression level of LC3 in the fractured femurs. Western blot was utilized to determine the expression level of PI3K, p-AKT1, AKT1, p-mTOR, mTOR, p-p70S6K, and p70S6K.

**Results:** The ALP activity and expression of RunX2 in fractured osteoblasts were significantly suppressed by 3-methyladenine and elevated by rapamycin, 60, and 80 μM β-ecdysterone. The apoptotic state of fractured osteoblasts was enhanced by 3-methyladenine and alleviated by rapamycin, 60, and 80 μM β-ecdysterone. The state of autophagy both in fractured osteoblasts and femurs was inhibited by 3-methyladenine and facilitated by rapamycin and β-ecdysterone. Compared to control, Garrett score in 3-methyladenine group was significantly decreased and promoted in rapamycin and β-ecdysterone groups, accompanied by ameliorated pathological state. Lastly, the PI3K/AKT/mTOR pathway both in fractured osteoblasts and femurs was activated by 3-methyladenine and inhibited by rapamycin and β-ecdysterone.

**Conclusion:** β-ecdysterone might facilitate fracture healing by activating autophagy through suppressing PI3K/AKT/mTOR signal pathway.

## Introduction

Fracture healing is a complicated and well-organized regulatory progress induced by series of histological and biochemical changes. The osteocytes, inflammatory cells, blood supply, and cytokines surrounding the fracture site play an important role in the whole process of fracture healing [1]. Although osseous tissues are equipped with powerful self-healing capacity and currently the therapeutic theoretical system against fracture healing is targeted and specific, approximately 5%-10% patients suffer from defective condition during the processing of fracture healing [2]. Currently, in the clinic, delayed fracture union or nonunion is commonly observed after treatments on fracture, which brought great difficulty and unnecessary burden for the patients [3]. Therefore, the quality of fracture healing is closely related to the living condition of the patients after treatments.

Autophagy is an important cellular mechanism maintaining the balance between cellular survival and cell death under stress state [4]. It is reported that autophagy is involved in the pathogenesis of multiple types of diseases, such as cancer, cardiac failure, hypertension, diabetes, and nerve diseases, by which several feasible strategies have been proposed for the treatment of these diseases [5, 6]. Recently, researches explored the impact of autophagy on the activity of osteoblasts by establishing animal models. Liu [7] claimed that the formation of autophagosome could be suppressed by downregulating the expression level of protein Fip200 in rat osteoblasts. As a consequence, the ossification in rats was significantly declined, which contributes to the poor bone growth and reduced sclerotin. By high throughput screening technology, rapamycin, an inducer of autophagy, is found to facilitate the differentiation and growth of osteoblasts [8]. These reports indicate that autophagy might exert an important role in the progress of bone growth and mineralization of bone tissues, which provides a potential therapeutic idea for the treatment of clinical delayed fracture union or nonunion.

β-ecdysterone is the main component of achyranthes bidentata and cyanotis arachnoidea, which is reported to exert multiple types of biofunctions, such as stimulating the synthesis of proteins, facilitating the metabolism of carbohydrates and lipids, alleviating hyperglycemia and hyperlipidemia, enhancing immunoregulation, and protecting the endothelial cells from apoptosis [9]. Gao reported that β-ecdysterone induced the in vitro differentiation from mesenchymal stem cells to osteoblasts in an estrogen receptor dependent manner [10]. Recently, it is reported that the autophagy of chondrocytes could be activated by β-ecdysterone to alleviate osteoarthritis [11]. In the present study, the protective effects of β-ecdysterone on fracture animal model, as well as the underlying mechanism, will be investigated to explore the potential therapeutic effect of β-ecdysterone on clinical delayed fracture union or nonunion.

## Materials and methods

### The establishment of fracture model in rats

The unilateral fracture was conducted in the middle third of right femur of rats according to the principle described previously [12]. Firstly, the animals were anaesthetized using an intraperitoneal administration of 25 mg/mL ketamine hydrochloride (Ketolar, Barcelona, Spain). The femur was inserted with an intramedullary Kirschner wire 1 mm in diameter before the fraction model was conducted. The knee was conducted with an anterior approach to lateralis the patella and expose the two femoral condyles, followed by inserting the wire through the intercondylar line till the major trochanter without drilling. Subsequently, the wire was cut under the cartilaginous surface. A small bent was made on the wire and the incision was sutured with 3/0 silk and 3/0 absorbable sutures. Finally, the fracture was closed immediately. The animals were sacrificed with euthanasia 4 weeks later and the fractured femur was isolated.

### The isolation of primary osteoblasts from fracture rat model

After the animals were sacrificed with euthanasia, the skin was pull down and middle third of right femur was exposed, followed by removing periosteum and blood vessels and interstitial cartilage attached to the bone. 0.25% trypsin containing 0.02% EDTA was added to the bone chips to be incubated for 25 min following excising the bone tissues into 1-2 mm^3^ pieces at 37 °C. Subsequently, the bone chips were placed in 5 ml Hanks solution containing 0.1% (wt/vol) Collagenase I, 0.05% trypsin, and 0.004% EDTA, which were further shaken in a shaking incubator at 37 °C at a shaking speed of 200 rpm for 1 hour. The cells were collected and the cells were centrifugated at 1000 rpm for 8 min, which were further resuspended in α-MEM medium containing 10% fetal bovine serum. The isolated osteoblasts were incubated at 37□ and CO_2_ for the subsequent experiments.

### In vitro grouping

The in vitro experiments were divided into 6 groups. In control group, the osteoblasts from fracture rat model were incubated with blank α-MEM medium. The osteoblasts in the rapamycin group were treated with 100 nM rapamycin and the osteoblasts in the 3-methyladenine group were introduced with 10 mM 3-methyladenine. In β-ecdysterone groups, the osteoblasts from fracture rat model were treated with 40, 60, and 80 μM β-ecdysterone, respectively. The osteoblasts were harvested for subsequent experiments after being incubated for 24 hours.

### In vivo grouping

The in vitro experiments were divided into 6 groups. The fracture rats in the control group were intraperitoneally injected with normal saline. The animals in the rapamycin group were intraperitoneally administered with 1 mg/kg rapamycin and the rats in the 3-methyladenine group were intraperitoneally administered with 30 mg/kg 3-methyladenine. Lastly, the fracture rats in β-ecdysterone groups were intraperitoneally injected with 0.6 mg/kg, 0.8 mg/kg, and 1.0 mg/kg β-ecdysterone, respectively. Animals in all groups were dosed twice a week for a consecutive 4 weeks. Afterwards, the animals were sacrificed with euthanasia and the fractured femur was isolated.

### MTT assay

The optimized concentration of rapamycin, 3-methyladenine, and β-ecdysterone was determined by MTT assay in osteoblasts. Briefly, the osteoblasts were planted on plates (Corning, NY, USA) at 37°C for 24 h, followed by adding with rapamycin (10, 20, 40, 60, 80, 100, 120, 140 nM), 3-methyladenine (1, 2, 4, 6, 8, 10, 12, 14 mM) and β-ecdysterone (1, 2, 5, 10, 20, 40, 60, 80, 100 μM) for 24 h, 48h and 72 h, respectively. Subsequently, the medium was added with 10 μL of 5 mg/mL MTT solution (Roche, Basel, Switzerland), followed by adding the formazan diluted in about 200 μL of dimethylsulfoxide (Genview, Beijing, China) four hours later. A microplate reader (Roche, Basel, Switzerland) was used to detect OD values at 490 nm. The value (ODcontrol-ODtreatment)/ODcontrol was used to represent suppressive rate.

### Alkaline phosphatase (ALP) activity assay

Briefly, treated osteoblasts were planted on 12-well plates at a density of 5 × 10^4^/well, followed by being washed and harvested. Subsequently, 100 μL lysis buffer (Beyotime, Shanghai, China) was used to lyse the cells, followed by measuring the ALP activity levels (U/ml) using a commercial kit (Nanjing Jiancheng Bioengineering Research Institute, Nanjing, China) according to the instruction of the manufacturer.

### Acridine orange (AO)/Propidium Iodide (PI) staining

The staining solution was prepared by mixing the 1 mg/mL acridine orange and 1 mg/mL Propidium Iodide (AO/PI) in 1 mL PBS buffer, which was added into each sample and incubated in the dark for 15 min. Lastly, the cells were flipped upside down into a sliding glass and imaged under a laser scanning confocal microscope (Olympus, Toyko, Japan).

### Flow cytometer assay

After treated with different strategies, the osteoblasts were harvested in 1.5 mL tubes, followed by adding 10 μL fluorescently-labeled Annexin V reagent and 5 μL PI reagent to be incubated at room temperature for 10 min. Subsequently, approximately 200 μL cells were added into the flow tube containing 2 mL PBS and tested by the flow cytometry (BD, New Jersey, USA). Three independent assays were performed.

### Reverse transcriptase-polymerase chain reaction (qRT-PCR)

Total RNA was collected from the cells using a RNA Extraction Kit (Thermo Fisher Scientific, Waltham, USA) in terms of the instructions of the manufacture. RNA extracted was quantified with a NanoDrop spectrophotometer (Thermo Fisher Scientific, Waltham, USA). A specific RT primer was used to reverse-transcribe the complementary DNA. SYBR Premix Ex TaqTM (Thermo Fisher Scientific, Waltham, USA) with an Applied Bio-Rad CFX96 Sequence Detection system (Genscript, NanJing, China) was used in the real-time PCR procedure. The expression level of *Atg7* and *LC-3* was determined by the threshold cycle (Ct), and relative expression levels were calculated by the 2^-ΔΔCt^ method after normalization with reference to the expression of U6 small nuclear RNA. The expression level of GAPDH in the tissue was taken as negative control. Three independent assays were performed. The primers were shown in Table 1.

### Western blotting assay

Total proteins were isolated from cells and tissues using the Nuclear and Cytoplasmic Protein Extraction Kit (Thermo Fisher Scientific, Waltham, USA). Approximately 40 μg of protein was separated on 12% SDS-polyacrylamide gel (SDS-PAGE) and the gel was transferred to polyvinylidene difluoride (PVDF) membrane (Millipore, MIT, USA). The membrane was blocked with 5% nonfat dry milk in TBST (Trisbuffered saline/0.1% Tween-20, pH 7.4) for 1 h at room temperature and incubated overnight with primary rabbit anti-human antibodies to ATG7 (1:1000, CST, Boston, USA), LC3-I/LC3-II (1:1000, CST, Boston, USA), PI3K (1:1000, CST, Boston, USA), p-AKT1 (1:1000, CST, Boston, USA), AKT1 (1:1000, CST, Boston, USA), p-mTOR (1:1000, CST, Boston, USA), mTOR (1:1000, CST, Boston, USA), p-p70S6K (1:1000, CST, Boston, USA), p70S6K (1:1000, CST, Boston, USA), and GAPDH (1:1000, CST, Boston, USA). A horseradish peroxidase-conjugated antibody against rabbit IgG (1:5000, CST, Boston, USA) was used as a secondary antibody. Blots were incubated with the ECL reagents (Amersham Pharmacia Biotech, Inc, USA) and exposed to Tanon 5200-multi to detect protein expression. Three independent assays were performed.

### Qualitative analysis of bone callus examination by X-ray

After all the treatments, X-ray was performed on the animals. All the animals were in supine position, with right femur flexed at a 90° angle. The exposure conditions were settled as 45KV, 0.08 s interval, and 70 mA. The state of fracture healing was evaluated by the Garrett scores on the X-ray results, which was determined by three senior orthopedists according to the standards of Garrett scores shown in Table 2.

### HE staining

After all the treatments, the calvaria and femurs were collected following sacrificing the animals. The liquid nitrogen was used to freeze the calvaria specimens and was fixed in 4% paraformaldehyde, followed by being dipped in 75% ethanol. Subsequently, the femurs were decalcified in 10% EDTA, followed by being embedded in paraffin and cut into sections at a 6 μm thick, which were further stained with hematoxylin and eosin (HE). Lastly, the sections were observed using a light microscope (Nikon, Tokyo, Japan) to evaluate the pathological changes of bone tissues.

### Immunohistochemical assay

In brief, the sections were deparaffinized and the ethylenediaminetetraacetic acid was used to perform the antigen retrieval in a pressure cooker, followed by incubating with hydrogen peroxidase. Subsequently, sections were incubated with primary anti-Beclin-1 antibody (Abcam, MA, USA), followed by being incubated with a secondary antibody (Abcam, MA, USA). The streptavidin labeled with horse radish peroxidase and 3,3’-Diaminobenzidine was used to perform the reaction. Lastly, the images were taken using a fluorescence microscope (Olympus, Toyko, Japan)

### Immunofluorescence assay

Briefly, sections were deparaffinated by xylene and washed twice using ethanol to remove remining xylene, which were dehydrated in the 95%, 80%, and 70% ethanol solution successively. Subsequently, sections were incubated with EDTA for 20 min, followed by being washed using PBS buffer and blocked by 3% goat serum for 1 hour at room temperature. Then, the samples were incubated with primary antibody against LC3-II at 4 □ overnight, followed by being incubated with FITC labeled anti-rat secondary antibody at 37 □ for 30 min. Finally, sections were sealed by neutral resins and observed under laser scanning confocal microscope (Olympus, Tokyo, Japan).

### Statistical analysis

Means ± standard deviation (SD) was displayed to show the data. Graphpad was used to analyze the data. Students’ t-test and one-way analysis of variance (ANOVA) were utilized for the contrast among different groups. *P*<0.05 was regarded as statistically significant difference between two groups.

## Results

### The determination of the concentrations of rapamycin, 3-methyladenine, and β-ecdysterone

To determine the optimized incubation concentration of drugs, MTT assay was conducted after the osteoblasts were incubated with different concentrations of rapamycin, 3-methyladenine, and β-ecdysterone. As shown in Fig 1, no significant difference on cell viability of osteoblasts was observed as the concentration of rapamycin increased from 10 nM to 100 nM, that of 3-methyladenine increased from 1 mM to 10 mM, and that of β-ecdysterone increased from 1 μM to 80 μM, respectively. However, the cell viability declined greatly as the concentration of rapamycin was higher than 120 nM, that of 3-methyladenine was higher than 12 mM, and that of β-ecdysterone was higher than 100 μM, respectively. Therefore, 100 nM rapamycin, 10 mM 3-methyladenine, 40, 60, and 80 μM β-ecdysterone were used in the subsequent in vitro experiments.

**Fig 1.**
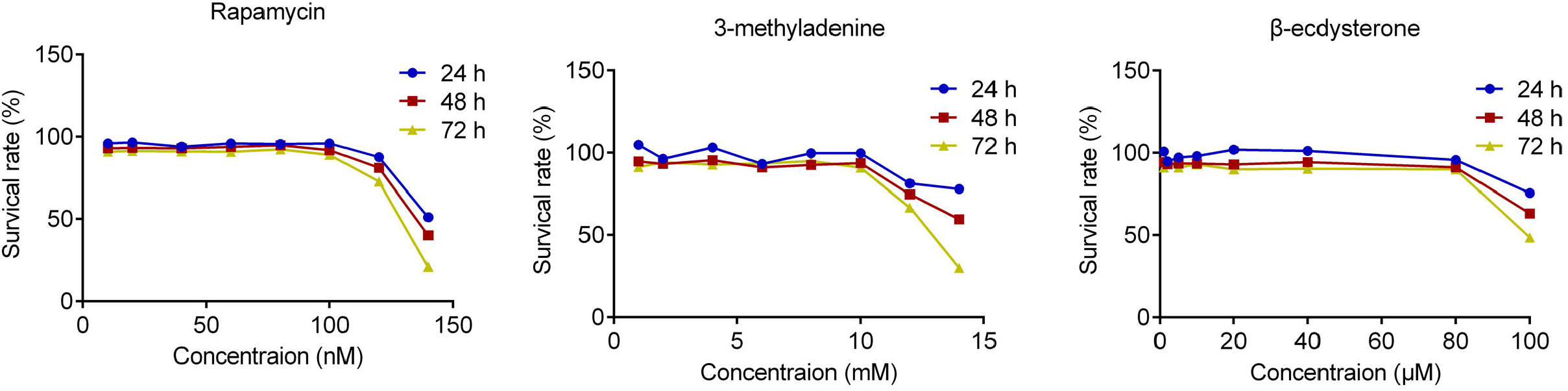
The cell viability at of osteoblasts treated with different strategies was evaluated by MTT assay.

### β-ecdysterone ameliorated osteogenic differentiation in osteoblasts

We further investigated the state of osteogenic differentiation in osteoblasts by detecting the ALP activity and the expression of RunX2. As shown in Fig 2A, the ALP activity in isolated osteoblasts was significantly suppressed by the incubation of 3-methyladenine and was dramatically promoted by the introduction of rapamycin, 60, and 80 μM β-ecdysterone, respectively. In addition, as shown in Fig 2B-C, compared to control, the gene and protein expression level of RunX2 was significantly inhibited by 3-methyladenine and was greatly elevated by the treatment of rapamycin, 60, and 80 μM β-ecdysterone, respectively (*p<0.05 vs. control, **p<0.01 vs. control).

**Fig 2.**
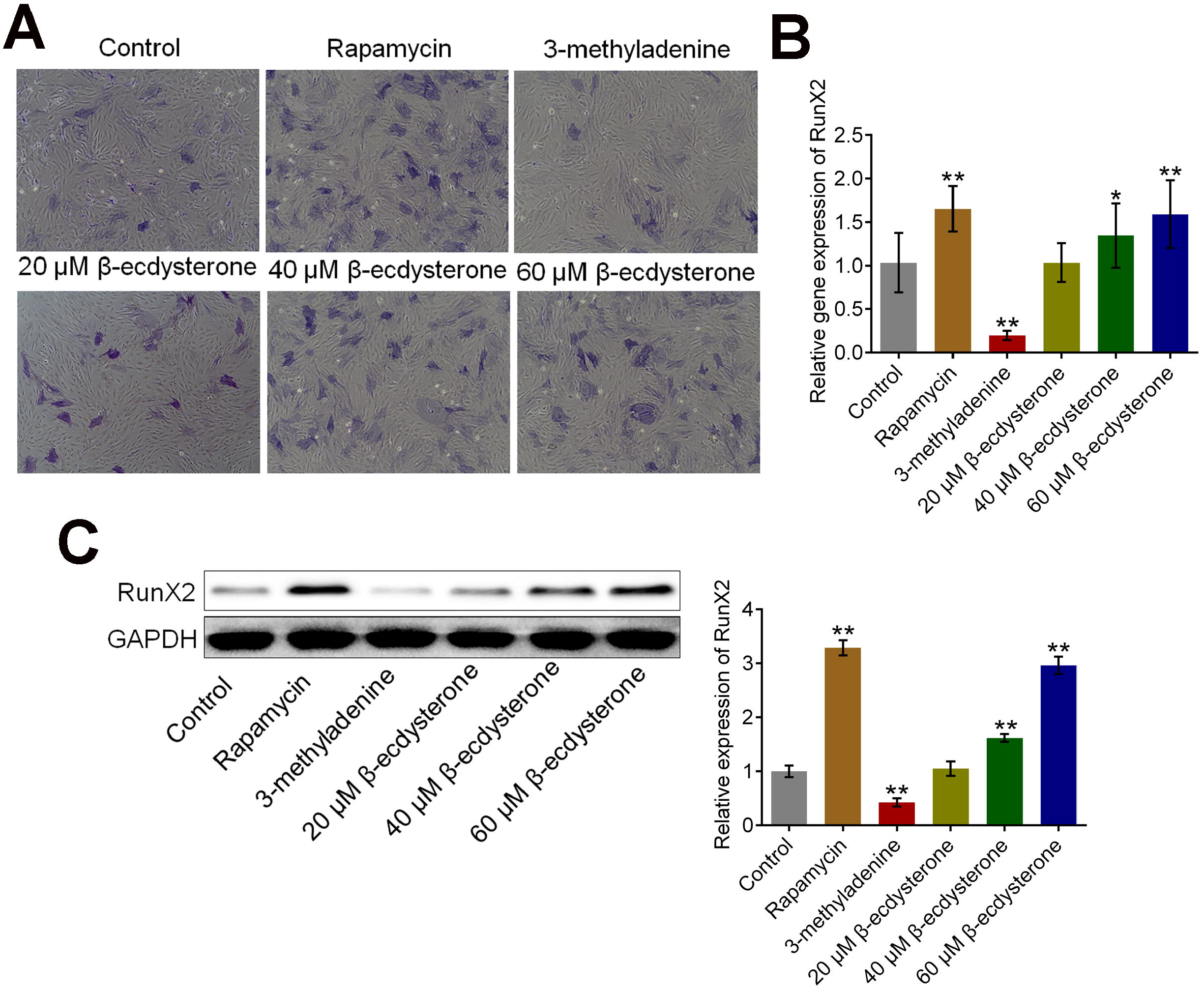
The osteogenesis differentiation in fractured osteoblasts was facilitated by β-ecdysterone. A. The ALP activity in each group was measured by a commercial kit. B. The gene expression level of RunX2 was detected by qRT-PCR assay. C. The expression level of RunX2 was determined by Western botting assay (*p<0.05 vs. control, **p<0.01 vs. control).

### β-ecdysterone alleviated the apoptosis of osteoblasts isolated from fracture rats

To explore the effects of β-ecdysterone on the apoptotic state of osteoblasts isolated from fracture rats, the AO/PI staining and flow cytometry assays were performed after the osteoblasts were treated with different strategies. As shown in Fig 3A, compared to control, the intensity of green fluorescence in 3-methyladenine group was significantly elevated and was dramatically suppressed in rapamycin, 60, and 80 μM β-ecdysterone groups, respectively. In addition, according to the results of flow cytometry assay, the apoptotic rate in the control, rapamycin, 3-methyladenine, 40, 60, and 80 μM β-ecdysterone groups was 26.62%, 17.04%, 34.4%, 25.74%, 20.64%, and 8.40%, respectively. These data indicated that the apoptotic state in osteoblasts isolated from fracture rats was significantly ameliorated by β-ecdysterone.

**Fig 3.**
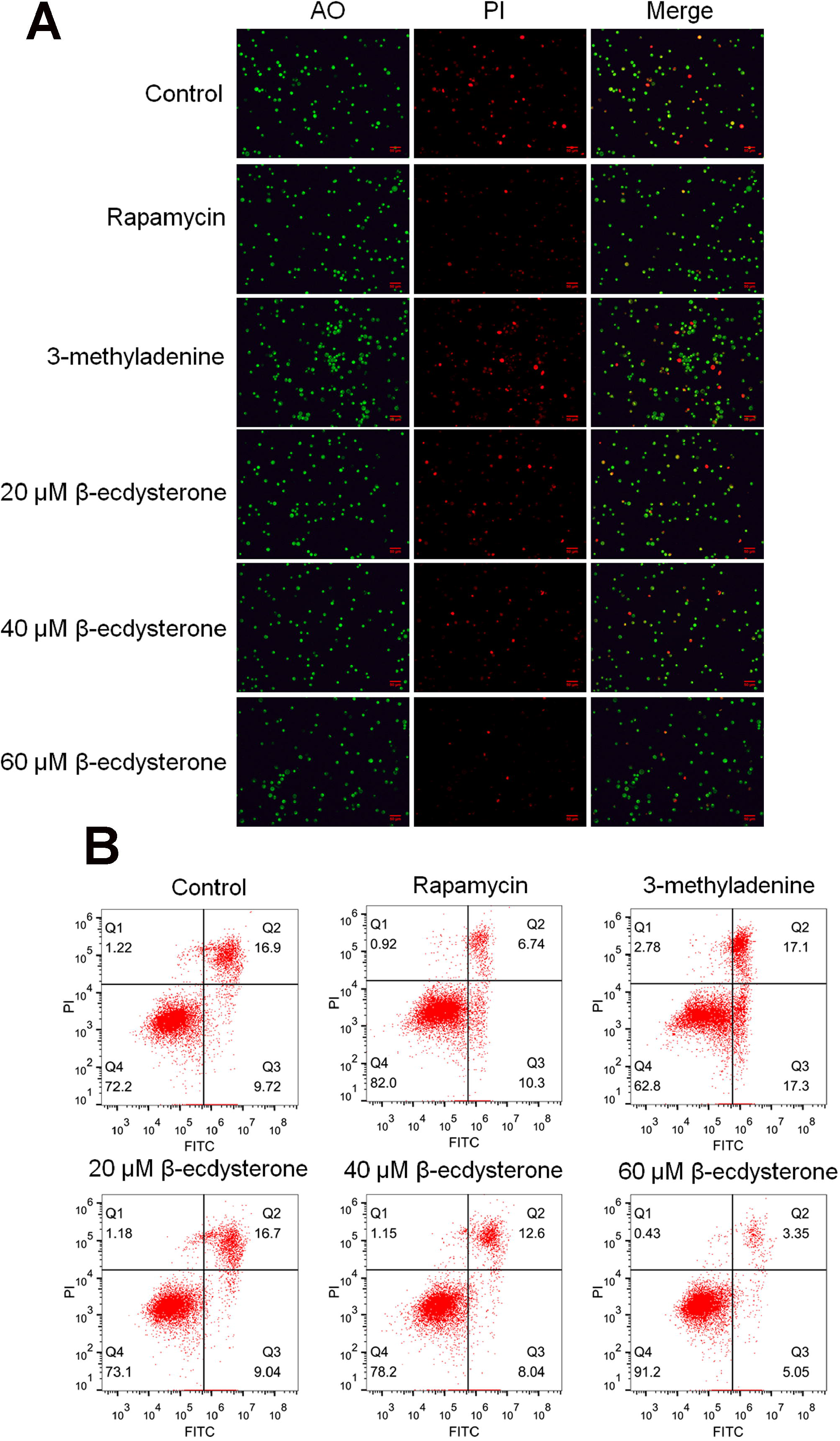
The apoptotic state of fractured osteoblasts was significantly alleviated by β-ecdysterone. A. The apoptotic state of treated osteoblasts was visualized by AO/PI staining. B. The apoptotic state of treated osteoblasts was quantified by flow cytometry assay.

### β-ecdysterone activated the state of autophagy in osteoblasts isolated from fracture rats

To explore the effects of β-ecdysterone on the state of autophagy in osteoblasts, the expression of ATG7 and LC3 was detected after osteoblasts were treated with different strategies. As shown in Fig 4, the expression level of both ATG7 and LC3-II/LC3-I was significantly suppressed by the introduction of 3-methyladenine and was dramatically elevated by the treatment of rapamycin, 60, and 80 μM β-ecdysterone, respectively (*p<0.05 vs. control, **p<0.01 vs. control). These data indicated that the state of autophagy in osteoblasts isolated from fracture rats as significantly activated by β-ecdysterone.

**Fig 4.**
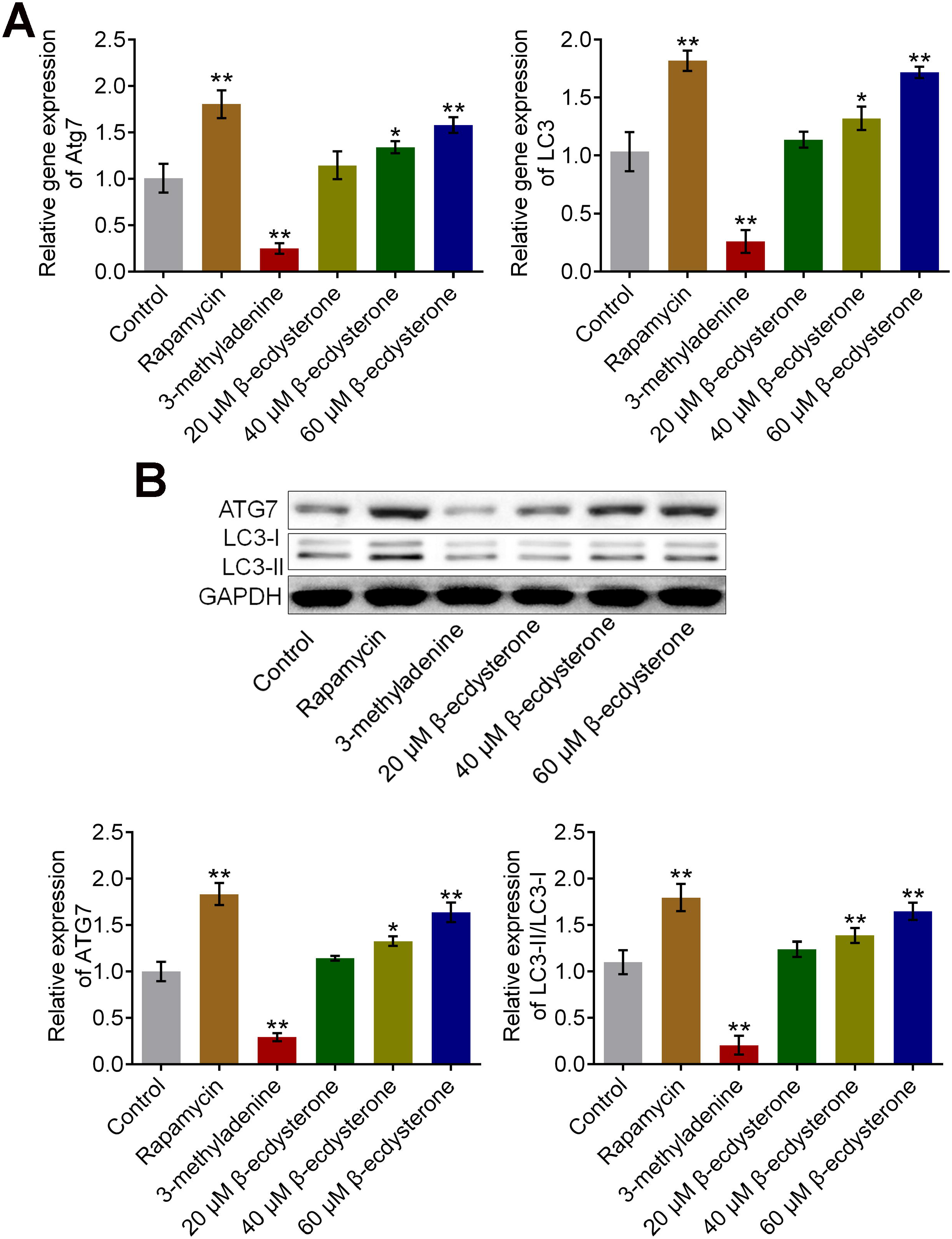
The autophagy in fractured osteoblasts was activated by β-ecdysterone. A. The gene expression level of ATG7 and LC3 was evaluated by qRT-PCR assay. B. The protein expression level of ATG7 and LC3 was determined by Western botting assay (*p<0.05 vs. control, **p<0.01 vs. control).

### β-ecdysterone inhibited the PI3K/AKT/mTOR signal pathway in osteoblasts isolated from fracture rats

We further investigated the expression of related proteins in PI3K/AKT/mTOR signal pathway, an important signal pathway that regulates autophagy, to explore the potential mechanism. As shown in Fig 5, PI3K, p-AKT1, p-mTOR, and p-p70S6K were significantly up-regulated in 3-methyladenine treated osteoblasts and were greatly down-regulated in rapamycin, 60, and 80 μM β-ecdysterone treated osteoblasts, respectively, indicating an inhibitory effect of β-ecdysterone on PI3K/AKT/mTOR signal pathway (*p<0.05 vs. control, **p<0.01 vs. control).

**Fig 5.**
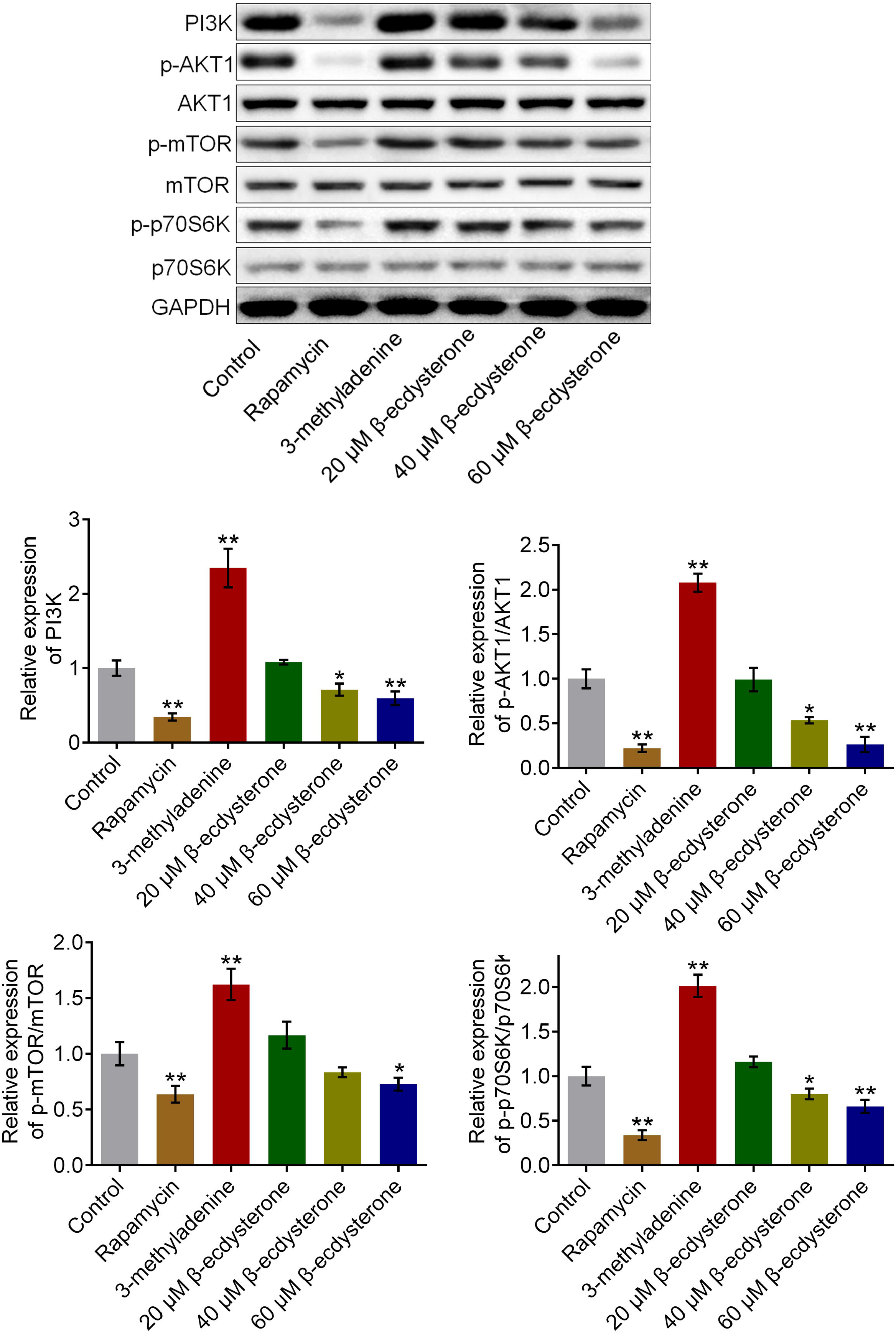
The PI3K/AKT/mTOR signal pathway in fractured osteoblasts was inhibited by β-ecdysterone. The expression level of PI3K, p-AKT1, AKT1, p-mTOR, mTOR, p-p70S6K, and p70S6K was detected by Western botting assay (*p<0.05 vs. control, **p<0.01 vs. control).

### β-ecdysterone accelerated the fracture healing of fracture rats

X-ray assay and HE staining were used to evaluate the condition of fracture healing in fracture rats. As shown in Fig 6 A and Table 3, compared to control, the fracture score was declined from 1.67 to 1.33 in 3-methyladenine group and was elevated to 3.67, 2.33, 2.83, and 3.50 in the rapamycin, 0.6 mg/kg, 0.8 mg/kg, and 1.0 mg/kg β-ecdysterone groups (*p<0.05 vs. control, **p<0.01 vs. control). In addition, in the control and 3-methyladenine groups, the results of HE staining (Fig 6B) showed that the direction of bone trabeculae was not explicit, the activity of osteogenesis was low, and the thickness of bone trabeculae was relatively thin. In the rapamycin, 0.6 mg/kg, 0.8 mg/kg, and 1.0 mg/kg β-ecdysterone groups, part of cartilaginous callus was replaced by osseous callus, some mature bone trabeculae and significant mineralization were observed, and the trabeculae were arranged orderly in line with the direction of stress. These data indicated that the fracture healing of fracture rats was dramatically accelerated by β-ecdysterone.

**Fig 6.**
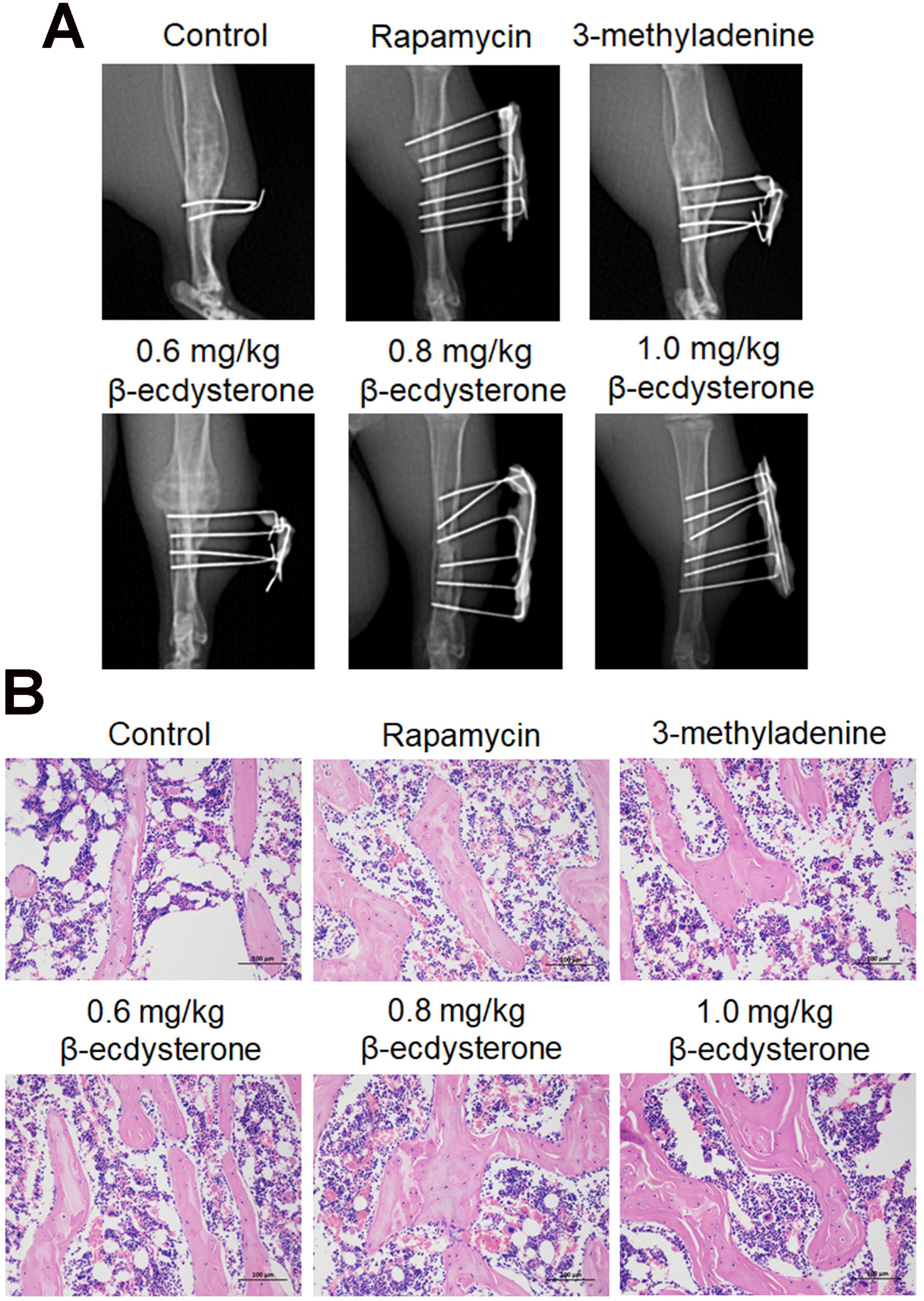
The pathological state in fractured femurs was dramatically alleviated by β-ecdysterone. A. X-ray assay was performed on fractured femurs isolated from each group. B. The pathological state in fractured femurs was visualized by HE staining assay.

### β-ecdysterone activated the state of autophagy in the fractured femur from fracture rats

After the animals were sacrificed, the fractured femurs were isolated and the autophagy was detected. The results of immunohistochemistry and immunofluorescence were shown in Fig 7A-B. compared to control, beclin-1 and LC3-II were significantly down-regulated in the 3-methyladenine group and were dramatically up-regulated in the rapamycin, 0.8 mg/kg, and 1.0 mg/kg β-ecdysterone groups. In addition, the results of Western blotting confirmed that compared to control, the expression of ATG7 and LC3-II/LC3-I was significantly inhibited in the 3-methyladenine group and was greatly elevated in the rapamycin, 0.8 mg/kg, and 1.0 mg/kg β-ecdysterone groups, respectively (*p<0.05 vs. control, **p<0.01 vs. control). These data indicated that the state of autophagy in the fractured femur from fracture rats was obviously activated by β-ecdysterone.

**Fig 7.**
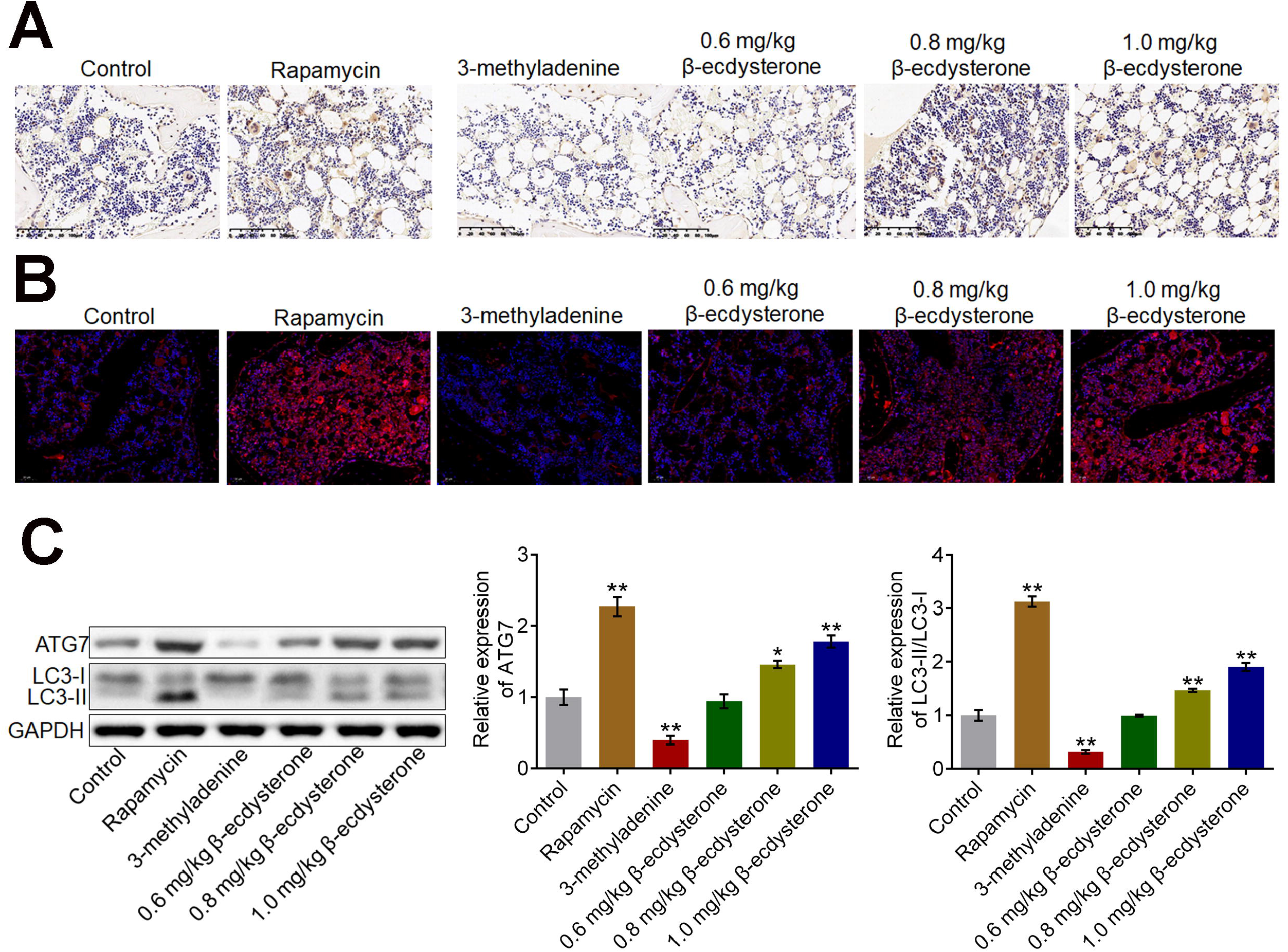
The autophagy in fractured femurs was activated by β-ecdysterone. A. The expression level of Beclin-1 in the fractured femurs was detected by immunohistochemical assay. B. The expression level of LC3 in the fractured femurs was determined by immunofluorescence assay. C. Western blotting assay was utilized to measure the expression level of ATG7 and LC3-II/LC3-I in the fractured femurs (*p<0.05 vs. control, **p<0.01 vs. control).

### β-ecdysterone inhibited the PI3K/AKT/mTOR signal pathway in the fractured femur from fracture rats

We further checked the state of PI3K/AKT/mTOR signal pathway in the fractured femur from fracture rats. As shown in Fig 8, the expression of PI3K, p-AKT1, p-mTOR, and p-p70S6K in the fractured femur from fracture rats was pronouncedly promoted in the 3-methyladenine group and was significantly suppressed in the rapamycin, 0.8 mg/kg, and 1.0 mg/kg β-ecdysterone groups, respectively (*p<0.05 vs. control, **p<0.01 vs. control), indicating that the PI3K/AKT/mTOR signal pathway in the fractured femur from fracture rats was obviously suppressed by β-ecdysterone.

**Fig 8.**
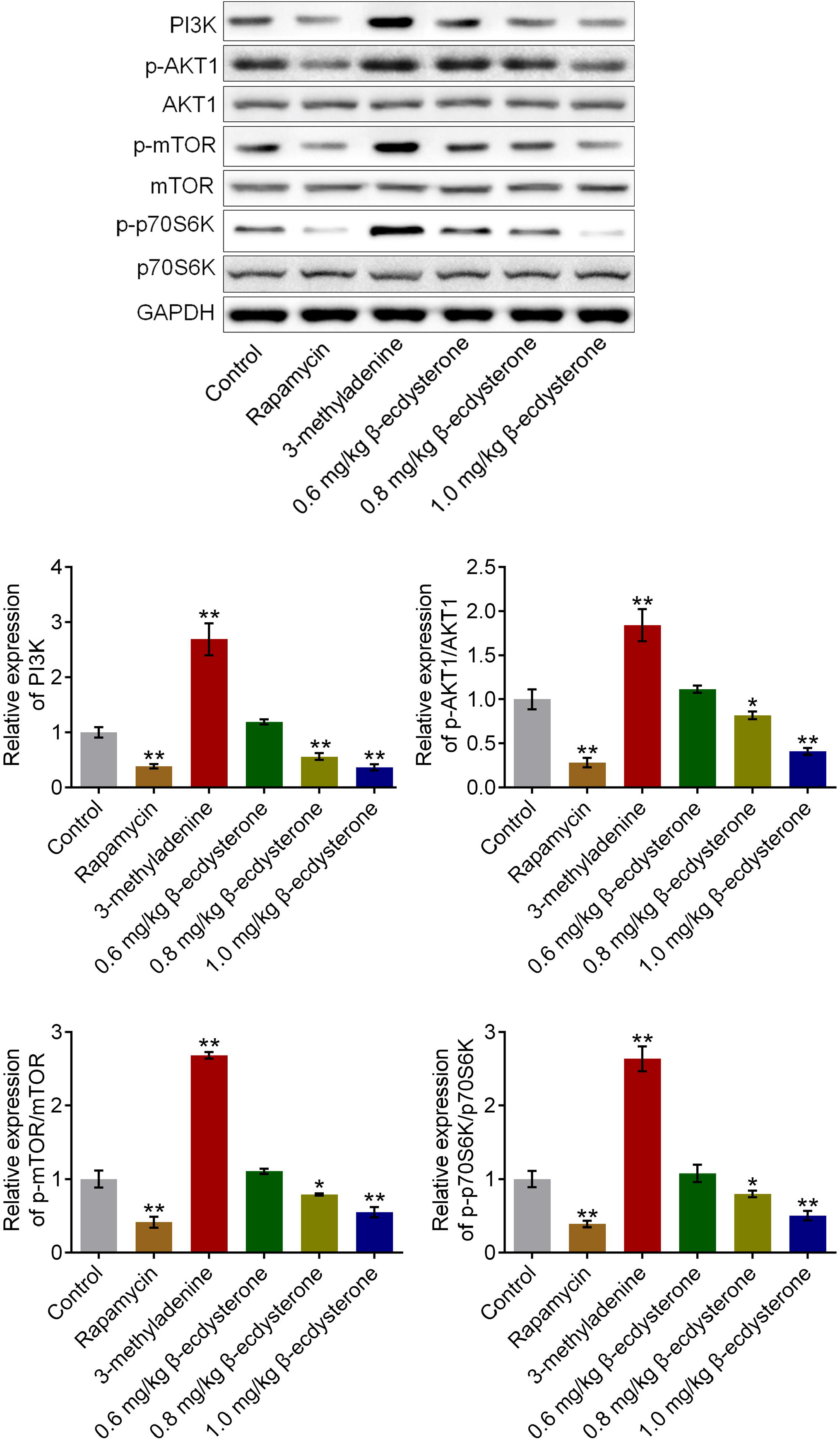
The PI3K/AKT/mTOR signal pathway in fractured femurs was inhibited by β-ecdysterone. The expression level of PI3K, p-AKT1, AKT1, p-mTOR, mTOR, p-p70S6K, and p70S6K was detected by Western botting assay (*p<0.05 vs. control, **p<0.01 vs. control).

## Discussion

Autophagy in osteoblasts has been widely reported to be involved in the mechanism of fracture healing. Li [13] reported that the femoral fracture healing in rat model was significantly promoted by curcumin by activating autophagy. Qiao [14] reported that the fracture healing in a rat model was restrained by inhibiting HIF-1α through suppressing autophagy. Therefore, the activation of autophagy contributes to the progression of fracture healing. The activation of autophagy is closely related to autophagy-related genes (ATG) [15] and microtubule associated protein 1 light chain 3 (LC-3) is an important regulator in the generation of autophagosome [16]. LC3 is located on the membrane of autophagy bubble and is divided into LC3-I and LC3-II. LC3-I is regularly expressed on the cellular membrane both in autophagy state and non-autophagy state. LC3-II is reported to be closely related to the development and processing of autophagy, which is generated from the binding between phosphatidyl ethanolamine on the membrane of autophagosome and the ubiquitin modified LC3-I. The cellular expression level of LC3-II is proportionally related to the number of autophagosome within the cells [17]. Beclin-1 is another biomarker of autophagy that is reported to regulate the activation of cellular autophagy in multiple types of cells [18, 19]. In the present study, we found that the osteogenic differentiation in osteoblasts and the fracture healing of fracture rats were significantly facilitated by the autophagy inducer rapamycin which was consistent with the reports described previously [20, 21]. We further verified that the osteogenic differentiation in osteoblasts and the fracture healing of fracture rats were significantly suppressed by the autophagy inhibitor 3-methyladenine. By the treatment of β-ecdysterone, the autophagy in both osteoblasts and the fractured femur was significantly activated, which was consistent with our previous data in chondrocytes [11]. The activation of autophagy induced by β-ecdysterone was found to be accompanied by facilitated osteogenic differentiation in osteoblasts and the fracture healing, indicating that the effects of β-ecdysterone on fracture healing might be related to the activation of autophagy in osteoblasts.

As an important cellular signal pathway, PI3K/AKT/mTOR pathway regulates cell apoptosis, autophagy, and cell proliferation by mediating the expression of downstream proteins [22]. In the PI3K/AKT/mTOR pathway, after AKT is activated by PI3K, the phosphorylation level of mTOR will be elevated, which finally contributes to the activation of autophagy [23]. In the present study, we found that the PI3K/AKT/mTOR pathway in both osteoblasts and the fractured femur was significantly suppressed by rapamycin and was greatly activated by 3-methyladenine, which was consistent with the reports claimed previously [24, 25]. By the treatment of β-ecdysterone, the PI3K/AKT/mTOR pathway in both osteoblasts and the fractured femur was dramatically suppressed, indicating that the effect of β-ecdysterone on autophagy might be related to the inactivation of PI3K/AKT/mTOR pathway.

Although we obtained preliminary data that supported the therapeutic effect of β-ecdysterone on the fracture healing, there were some limitation on the present study. Firstly, in the present study, osteoblasts isolated from fracture rats were used as control in the in vitro study and fracture rats was taken as control in the in vivo experiments, without involving the Sham group into the study design. In our future work, a Sham group will be introduced to further verify the successful establishment of fracture animal model. Although in the present study, the results of X-ray and HE staining assays provided sufficient evidence for the successful establishment of fracture rats. Secondary, we revealed that the effects of β-ecdysterone osteogenic differentiation and fracture healing were accompanied by the activation of autophagy and inhibition of PI3K/AKT/mTOR pathway. However, further verifications should be performed. In our future, the combination administration of β-ecdysterone and 3-methyladenine will be introduced in both in vitro and in vivo experiments to verify the involvement of autophagy and PI3K/AKT/mTOR pathway in the functional mechanism of β-ecdysterone on fracture healing.

Taken together, our data indicated that β-ecdysterone might facilitate fracture healing by activating autophagy through suppressing PI3K/AKT/mTOR signal pathway.

## Acknowledgments

This work was supported by grants from the Medical and Health Science and Technology Program of Zhejiang Province (Grant No. 2019KY548) and the Health Science and Technology Project of Hangzhou (Grant No. 2018B029).

